# Decreased kinesin-1 mitigates NMDA-induced exicitotoxicity and ischemia-evoked neurodegeneration

**DOI:** 10.1101/440800

**Authors:** Raozhou Lin, Zhigang Duan, Haitao Sun, Man-Lung Fung, Hansen Chen, Jing Wang, Chi-Fai Lau, Di Yang, Yu Liu, Yanxiang Ni, Zai Wang, Ju Cui, Wutian Wu, Wing-Ho Yung, Ying-Shing Chan, Amy C. Y. Lo, Jun Xia, Jiangang Shen, Jian-Dong Huang

## Abstract

N-methyl-D-aspartate receptor (NMDAR) is highly compartmentalized in neurons and the dysfunction has been implicated in various neuropsychiatric and neurodegenerative disorders. Recent failure to exploit NMDAR antagonization as a potential therapeutic target has driven the need to identify molecular mechanisms that regulate NMDAR compartmentalization. Here, we report that neural activity-dependent reduction of Kif5b, the heavy chain of kinesin-1, protected neurons against NMDA-induced excitotoxicity and ischemia-provoked neurodegeneration. Direct binding of Kinesin-1 to the GluN2B cytoplasmic tails regulated levels of NMDAR at extrasynaptic sites and the subsequent influx of calcium mediated by extrasynaptic NMDAR via regulating the insertion of NMDARs into neuronal surface. Transient increase of Kif5b restored the surface levels of NMDAR and the decreased neuronal susceptibility to NMDA-induced excitotoxicity. Our findings reveal that kinesin-1 regulates extrasynaptic NMDAR targeting and signaling, and the reduction of kinesin-1 could be regulated by neural activity and could be exploited to postpone or halt neurodegeneration.

## Introduction

The prevalence of neuropsychiatric and neurodegenerative disorders is increasing and is putting increasing pressure on health systems. Despite differences in the pathologies among various disorders, evidences suggest that N-Methyl-D-aspartic acid receptors (NMDARs) dysfunction is one of the common causes(Paoletti, Bellone et al., 2013, Zhou & Sheng, 2013). NMDARs belongs to a group of glutamate-gated ion channels that are essential for mediating brain plasticity(Paoletti et al., 2013). These receptors have been known for decades to receive input from presynaptic neurons and convey these signals into different patterns of neuronal activity, which eventually leads to long-term changes that underlies higher cognitive functions(Paoletti et al., 2013). Pathophysiologically, overactivation of NMDARs activates subsequent pro-death pathways such as calcium influx and the production of reactive oxygen species, which are the main causes of synaptic dysfunction and neuronal death underlying neurodegenerative disorders, like stroke and Alzheimer’s diseases. Recently, ketamine, an NMDAR antagonist, has been found to elicit rapid antidepressant effect(Murrough, Abdallah et al., 2017). Despite huge clinic potential, current attempts to translate NMDAR blockade into clinical applications have so far been unsatisfactory. The concern raised in inhibiting NMDARs might result from the complexity of NMDAR compartmentalization.

There are two distinct pools of NMDARs. Comparing to the well-established role of the synaptic NMDARs in mediating synaptic plasticity(Paoletti et al., 2013, Thomas & Huganir, 2004), the understanding on the extrasynaptic NMDARs is elusive. It has been reported that extrasynaptic NMDAR is activated by excessively released and spillovered glutamate, and consequently shuts off the pro-survival pathway elicited by synaptic NMDARs(Bading, 2013, Hardingham, Fukunaga et al., 2002). The extrasynaptic NMDARs induced excitotoxicity has been implicated in various neurodegenerative conditions, including stroke and Alzheimer’s disease(Bading, 2017, Molokanova, Akhtar et al., 2014, Okamoto, Pouladi et al., 2009, Sattler, Xiong et al., 1999, Tu, Xu et al., 2010). As such, direct and overall inhibition of NMDARs might disrupt NMDAR signaling in all compartments, including the physiological NMDAR signaling that benefits the neurons(Bading, 2017, De Keyser, Sulter et al., 1999, Sanacora, Frye et al., 2017). It is therefore urged to understand the mechanism that regulates the compartmentalization of NMDARs, which can be exploited to abate pathophysiological NMDAR signaling with minimal adverse physiological impacts.

Due to their highly polarized structure, neurons rely heavily on intracellular transportation systems to maintain their the highly compartmentalized signaling that integrate and process information from different inputs and sub-neuronal compartments (Hirokawa, Niwa et al., 2010, Terenzio, Schiavo et al., 2017). Kinesin-1 is a molecular motor complex consisting of two heavy chains and two light chains. One of the heavy chains, Kif5b, is comprised of three functional domains: the head domain drives anterograde transport along microtubules powered by ATP hydrolysis, the stalk domain facilitates the dimerization of heavy chains, and the tail domain mediates cargo binding (Hirokawa et al., 2010, Verhey, Kaul et al., 2011). Kinesin-1 with intact Kif5b is critical for development, as demonstrated by homozygous knockout of Kif5b in mice, which causes embryonic death as early as 9.5 days *in utero* (Tanaka, Kanai et al., 1998). Moreover, Kinesin-1 dysfunction is implicated in the pathogenesis of various neurodegenerative disorders. For example, Kif5b-containing vast vesicles, an early hallmark of axonal transport defect, are found in the autopsy brain of Alzheimer’s disease (Stokin, Lillo et al., 2005), and reduced levels of kinesin heavy chain are found in the early stages of Parkinson’s disease that precedes alteration of dopaminergic markers (Chu, Morfini et al., 2012).

Here we identified kinesin-1 as a microtubule dependent molecular motor that regulates the distribution and function of extrasynaptic NMDARs. Kinesin-1 binds with GluN2B NMDAR via their carboxyl tails. The reduction of kinesin-1 prevents extrasynaptic NMDAR targeting, inhibits calcium influx from extrasynaptic NMDARs and protects neuron against NMDA elicited exicitotoxicity and ischemia evoked neurodegeneration. Furthermore, the expression of kinesin genes, including *kif5b*, is reduced in the transcriptome of cerebral ischemia preconditioning, indicating the reduction of kinesin-1 as an intrinsic neuroprotective mechanism via regulating the compartmentalization of signaling.

## Results

### Kif5b binds with GluN2B-containing NMDARs

To test the hypothesis that kinesin-1 regulates the compartmentalization of NMDARs, we firstly immunoprecipitated endogenous kinesin-1 from mouse brain lysate using antibodies against Kif5b and detected GluN1, GluN2A, and GluN2B NMDAR subunits in the precipitate (Fig. 1a). Reciprocal immunoprecipitation using an antibody against GluN 1 further confirmed the presence of this complex (Fig. S1a). Moreover, Kif5b and GluN2B were found to share partial co-localization in both soma and dendrites as seen in labeled primary hippocampal neurons at 14 days *in vitro* (DIV) using corresponding antibodies (Fig. 1b). This *in vivo* data revealed a novel protein complex consisting of kinesin-1 and NMDAR.

**Figure 1.**
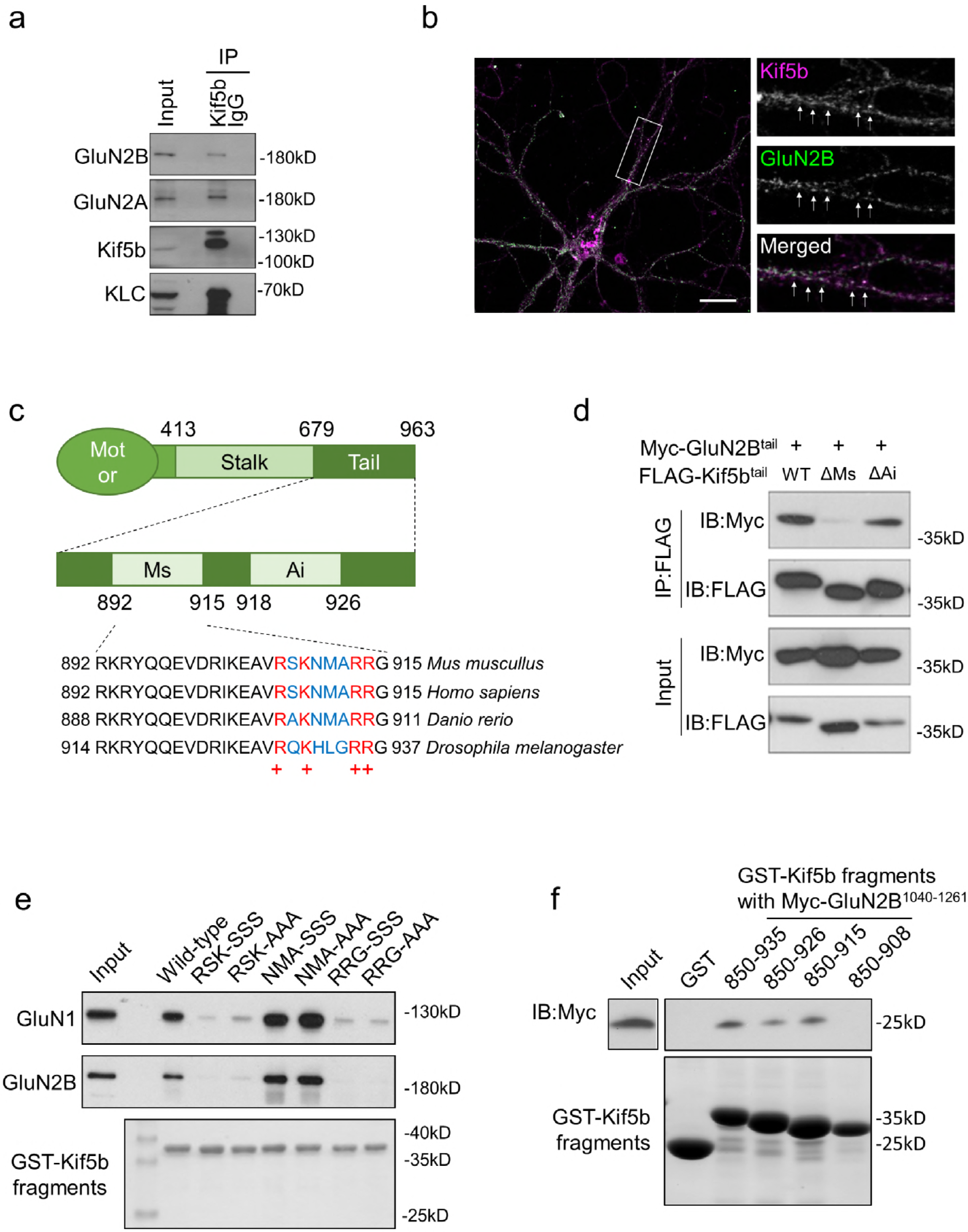
Kif5b directly binds with GluN2B. (a) Representative Western blot images of Kif5b or normal IgG immunoprecipitation from mouse brain lysate. (b) Immunostaining of Kif5b and GluN2B in primary hippocampal neurons (14 DIV). Scale bar (white solid line) = 10 μm. The arrows in the “Merge” panel indicate the co-localization signals. (c) Schematic figure of Kif5b fragment containing the motor domain (1-413 aa), the stalk domain (414-679 aa), the tail domain (679-963 aa), the microtubule sliding domain (892-915 aa), and the autoinhibitory domain (918-926 aa). The microtubule sliding domain is conserved from *Drosophila melanogaster* to *Homo Sapiens*. The four positively charged amino acids are shown in red. (d) Representative Western blot image of Kif5b tail immunoprecipitated with GluN2B intermediate tail (1040-1261 aa) from 293T cells overexpressing the indicated constructs. (e) Representative Western blot images of GST-tagged Kif5b tail fragment (900-935 aa) and its mutants with pull down of GluN1 and GluN2B from mouse brain lysate. (f) Representative Western blot image of GST-tagged Kif5b tail directly binding with in vitro transcribed and translated GluN2B intermediate tails.

The Kif5b protein consists of the head domain that acts as the motor for moving along the microtubule, the coiled-coil stalk domain that forms a motor complex with other heavy chains, and the tail domain that is believed to bind to cargo. Two functional sites within the tail domain have been identified, a microtubule sliding site and an autoinhibitory site(Kaan, Hackney et al., 2011, Wong & Rice, 2010). We generated a series of Kif5b truncations conjugated with a GST tag in a pull-down experiment to investigate this interaction in detail (Fig. 1c). We found the Kif5b tail (850-915 aa) including the microtubule sliding site, which was outside of the KLC binding domain, mediated binding to NMDAR (Fig. S1, b and c). This mapping result was confirmed by co-expression of FLAG-tagged Kif5b fragments and the intermediate tail (1040-1261 aa) of GluN2B in a 293T cell line and co-immunoprecipitation by FLAG (Fig. 1d). It is worth noted that deletion of the microtubule sliding domain (892-915 aa, ΔMs), but not the autoinhibitiory domain (918-926 aa, ΔAi), largely abolished these interactions, suggesting this domain was important in mediating Kif5b binding with NMDARs (Fig. 1d). By examining this interaction in further detail, we identified four positively charged amino acids that are conserved across species within this region (Fig. 1c) and wondered whether these positive charged amino acids are required for binding with NMDAR. Mutations of either of the two positively charged amino acids (907-909 aa, RSK to SSS or AAA; 913-915 aa, RRG to SSS or AAA) abolished this binding, whereas mutations of other amino acids (910-912 aa, NMA to SSS or AAA) did not cause any disruption (Fig. 1e). Furthermore, these were direct interactions, because GST-tagged Kif5b tail fragments were able to bind with in vitro transcribed and translated GluN2B intermediate tails in a cell-free system (Fig. 1f). Above data combined reveals a novel complex consisting of kinesin-1 and GluN2B-containing NMDARs, indicating a role of kinesin-1 in mediating the function of GluN2B-NMDARs.

### Kinesin-1 regulates extrasynaptic NMDAR distribution and function

Kinesin-1 regulates transportation of cargo along microtubules through direct binding to cargo or through various adaptor proteins. This motor-cargo interaction determines the subcellular distribution of cargo and thus regulates their functions(Hirokawa et al., 2010). It has been previously reported that GluN2B-containing NMDAR is conveyed by Kif17(Setou, Nakagawa et al., 2000, Yin, Takei et al., 2011). Interestingly, depletion of *kif17* did not completely abolish the movement of GluN2B *in vivo*(Yin et al., 2011), raising the possibility that multiple transport systems might contribute to regulate GluN2B-containing NMDAR trafficking. To examine the functional impact of kinesin-1 on GluN2B *in vivo*, we utilized a mouse model with heterozygous knockout of *kif5b* genes, as the homozygous knockout of kif5b causes embryonic lethality(Tanaka et al., 1998). As expected, besides the reduced levels of Kif5b, none of the other tested kinesins was found altered including Kif5a, Kif5c, Kif17, and Kif3 (Kif5b, *P*=0.0013, t=4.4294, df=10; Kif5a, *P*=0.5934, t=0.5535, df=9; Kif5c, *P*=0.2031, t=1.3619, df=10; Kif17, *P*=0.6209, t=0.5103, df=10; Kif3, *P*>0.9999, t=0, df=10; unpaired t-test; Fig. S2a). *Kif5b*^+/−^ hippocampus displayed reduced levels of the kinesin-1-GluN2B complex as revealed by immunoprecipitation of Kif5b (*P*=0.0005, t=5.603, df=8; unpaired t-test; Fig. S2b). Because the level of Kif17 was not altered in our *kif5b*^+/−^ mice (Kif17, *P*=0.6209, t=0.5103, df=10; unpaired t-test; Fig. S2a), it appears kinesin-1 reduction does not simultaneously affect Kif17-mediated transport. As such, we hypothesized that Kif5b might serve as a motor necessary for GluN2B-containing NMDAR trafficking. The Western blot analysis showed no differences in the overall levels of NMDAR subunits GluN1, GluN2A, and GluN2B in hippocampal lysate in wild-type and *kif5b*^+/−^ mice (GluN1, *P*=0.5405, t=0.6362, df=9; GluN2A, *P*=0.4666, t=0.7602, df=9; GluN2B, *P*=0.3506, t=0.9845, df=9; unpaired t-test; Fig. S2c). However, surface biotinylation of hippocampal slices from *kif5b*^+/−^ and control mice showed significantly reduced levels of surface GluN2B-containing NMDARs (GluN1, *P*=0.0005, t=4.672, df=12; GluN2A, *P*=0.5826, t=0.5727, df=8; GluN2B, *P*<0.0001, t=6.647, df=12; unpaired t-test; Fig. S2d). This was further confirmed by surface labeling of primary neurons transfected with GFP-tagged GluN2B, which showed both surface GFP puncta density and intensity were reduced in *kif5b*^+/−^ neurons at 14-16 DIV (Fig. 2, a to c; c, left, density, *P*<0.0001, D=0.6448; right, intensity, *P*<0.0001, D= 0.5961; Kolmogorov-Smirnov test). To test whether kinesin-1 affects dynamic turnover GluN2B-containing NMDAR, we performed a surface antibody labeling assay on neuronal culture from *kif5b*^+/−^ mice and their control littermates. We first saturated the GFP-GluN2B transfected neurons with anti-GFP antibody raised in chicken. After thorough wash, the neurons were returned to 37°C for additional 2 hrs to allow GluN2B turnover. The neurons were further incubated with anti-GFP antibody raised in rabbit to distinguish the original surface GluN2B from the newly inserted GluN2B (Fig. 2d). Compared with the that of the control littermates, there was significantly reduced amount of newly inserted GluN2B in the *kif5b*^+/−^ cultures (*P*<0.0001, U=22.5, Mann-Whitney U test; Fig. 2, e and f).

**Figure 2.**
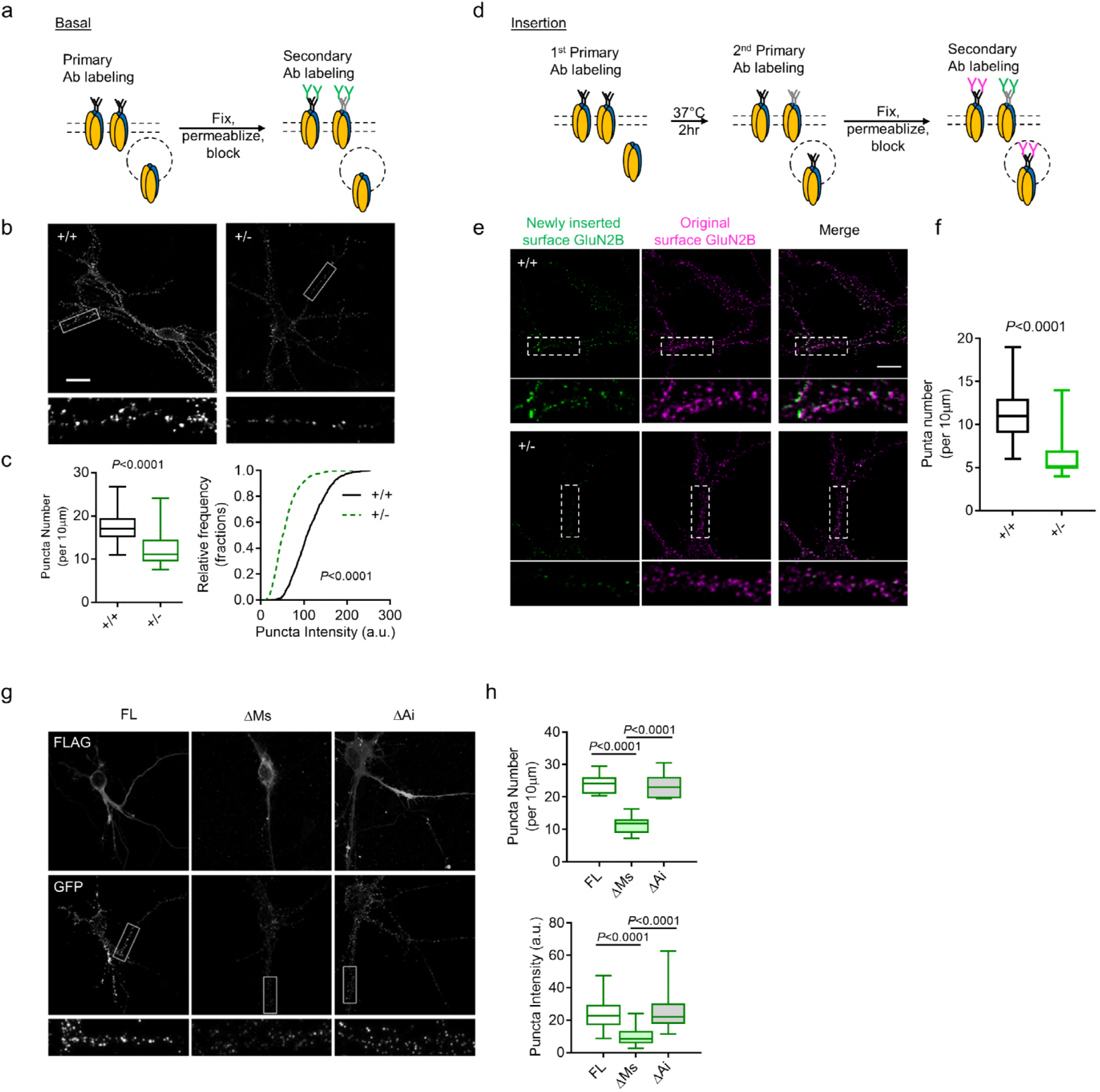
Kinesin-1 regulates NMDAR surface expression. (a) Schematic figure showing antibody labeling the surface GluN2B-containing NMDARs. Plasmid with extracellular GFP tagged GluN2B was transfected into primary neuronal cultures at 14 D. I. V‥ After 24 hrs, neurons were incubated with anti-GFP antibody at 4°C for 1 hr, followed by fixation and secondary antibody incubation. (b) Representative images and (c) quantification of surface GFP-GluN2B staining in primary neuronal cultures from *kif5b*^+/−^ mice (+/−) and their wild-type littermates (+/+). (c, left) Box plot showed the number of puncta per 10 μm while (c, right) the curve plots show the relative intensity of each puncta (puncta number: n=41 for wild-type and n=39 for *kif5b*^+/−^, five neurons were analyzed per genotype, Mann-Whitney U test; puncta intensity: six neurons were analyzed per genotype, at least 100 puncta were analyzed per neuron from three individual cultures, Kolmogorov-Smirnov test). Scale bar (white solid line) = 20 μm. (d) Schematic figure showing antibody labeling for the newly inserted GluN2B-containing NMDARs. After transfection, the neurons were saturated with chicken anti-GFP antibody for 10 min at room temperature and washed before returned to 37°C for additional 2 hrs. Then neurons were incubated with rabbit anti-GFP antibody for another 10 min at room temperature to label the newly inserted GluN2B. The neurons were washed, fixed and permeabilized for secondary antibody labeling to distinguish between the newly inserted and the original GluN2B. (e) Representative images and (f) quantification of surface GFP-GluN2B staining in primary neuronal cultures from *kif5b*^+/−^ mice (+/−) and their wild-type littermates (+/+). (f) Box plot showed the number of puncta of newly inserted GluN2B (green channel) per 10 μm. (n=15 per genotype; at least three neurons per genotype; for each neuron, at least three neurites were analyzed; Mann-Whitley U test). Scale bar (white solid line) = 10 μm. (g) Representative images show surface staining of GFP-GluN2B puncta from neurons co-transfected with the indicated Kif5b constructs. Bar plots show (h, left) the average density of surface puncta and (h, right) the intensity of each puncta (h, left: n=14 for FL, n=13 for ΔMs, n=19 for ΔAi, eight neurons analyzed for each group; h, right: n=167 for FL, n=128 for ΔMs, n=137 for ΔAi, Kruskal-Wallis test followed by Dunn’s *post hoc* test, 20 puncta/neuron analyzed for each group; three replicate experiments). Scale bar (white solid line) = 20 μm. Values are presented as mean ± s.e.m., unless otherwise indicated (test as indicated; exact *P* value given if less than 0.05).

We further examined the effect of the identified domain on mediating the GluN2B-containing NMDAR distribution. The 3×FLAG-tagged full-length Kif5b, Kif5b with microtubule sliding site deletion, or Kif5b with autoinhibitory site deletion together with GFP-GluN2B were transfected into primary cultures from *kifib*^+/−^ mice, respectively. Compared with neurons expressing Kif5b with microtubule sliding site deletion, both wild-type Kif5b and Kif5b with autoinhibitory site deletion showed increased levels of both GluN2B surface puncta density and puncta intensity (density, H=27.96, *P*<0.0001; FLvs. ΔMs, *P*<0.0001; FLvs. ΔAi, *P*=0.4242; ΔMs vs. ΔAi, *P*<0.0001; intensity, H=217.0, *P*<0.0001; FLvs. ΔMs, *P*<0.0001; FLvs. ΔAi, *P*=0.3565; ΔMs vs. ΔAi, *P*<0.0001; Kruskal-Wallis test followed by *post hoc* Dunn’s multiple comparisons test; Fig. 2, g and h).

NMDAR acts as a calcium channel upon activation. The resulting calcium influx regulates synaptic strength(Bading, 2013, Volk, Chiu et al., 2015), or causes the pro-death signaling cascade leading to neuronal death when it is overloaded(Bading, 2013, Tu et al., 2010). NMDAR-dependent calcium influx is associated with synaptic dysfunction and neuronal loss in both acute and chronic neurodegenerative conditions(Bading, 2013, Zhou & Sheng, 2013). To gain insight on the functional impact of kinesin-1 on NMDAR compartmentalization, we used Fluo-3 to quantify calcium influx. In *kifib*^+/−^ neurons, we found NMDA-induced calcium influx was significantly reduced (Genotype, F(1, 12000) = 17333, *P*<0.0001; twoway ANOVA followed by Sidak *post hoc* test; Peak, *P*<0.0001, U=68.00; Mann-Whitney U test; Fig. S3, a and b), possibly indicating NMDAR hypofunction or a more generalized defect in calcium homeostasis. To distinguish between these two possibilities, we depolarized the neurons with KCl (50 mM), which triggered a similar level of calcium influx in both genotypes (Genotype, F(1, 4240) = 19.63, *P*<0.0001; two-way ANOVA followed by Sidak *post hoc* test; Peak, *P*=0.1561, U=289.0; Mann-Whitney U test; Fig. S3c). This result reassured that the reduced calcium influx in *kif5b*^+/−^ was caused by NMDAR hypofunction.

On the neuron surface, NMDARs are distributed in postsynaptic densities (PSD) as well as in extrasynaptic regions, and the compartmentalization determines their signaling and functions. Given the notion that GluN2B is predominately concentrated in extrasynaptic regions in adult mouse hippocampus(Tovar & Westbrook, 1999), we hypothesized that the kinesin-1-NMDAR complex would biasedly regulates the targeting and functioning of extrasynaptic NMDARs. To test this hypothesis, we immunostained surface GFP-GluN2B together with the PSD marker, PSD-95, to distinguish different NMDAR compartments at the same dendrite. While similar numbers of synaptic puncta containing GluN2B and PSD-95 was found both wild-type and *kif5b*^+/−^ groups (*P*=0.3767, t=1.266, df=58; two-way ANOVA followed by Sidak *post hoc* test), we observed a significant reduction of surface GFP-GluN2B signals that were not co-localized with PSD-95 in *kif5b*^+/−^ neurons (*P*<0.0001, t=9.280, df=58; two-way ANOVA followed by Sidak *post hoc* test; Fig. 3, a and b). This data showed that the surface targeting of extrasynaptic NMDARs were dampened by reducing kinesin-1 level. To examine the functional impact of kinesin-1 on extrasynaptic NMDAR, we monitored the calcium influx elicited by bicuculline, which increase neuronal activity by blocking a GABAA receptor, together with 4-AP. We observed a calcium influx that was believed to be synapse-specific. We then added MK-801, which targets NMDAR calcium channel opening, to block only the activated NMDAR(Hardingham et al., 2002). We did not detect any notable differences in calcium influx between the wild-type and *kif5b*^+/−^ neurons, suggesting synaptic NMDAR-dependent calcium influx was not altered in *kif5b*^+/−^ neurons (*P*=0.9550, U=980.0; Mann-Whitney U test; Fig. 3e). As MK-801 is believed to irreversibly block extrasynaptic NMDAR, we re-stimulated the neurons after MK-801 blockade with NMDA to examine extrasynaptic NMDAR-dependent calcium influx, which showed a significant reduction in *kif5b*^+/−^ neurons (*P*<0.0001, U= 326.0; Mann-Whitney U test; Fig. 3, c to e). Taken together, the above data showed that kinesin-1 binding with GluN2B mediates NMDAR compartmentalization by regulating the extrasynaptic targeting of NMDARs, and thereby regulates NMDA-elicited calcium influx. This further indicates a role of kinesin in determine cargo destination towards different neuronal sub-compartments.

**Figure 3.**
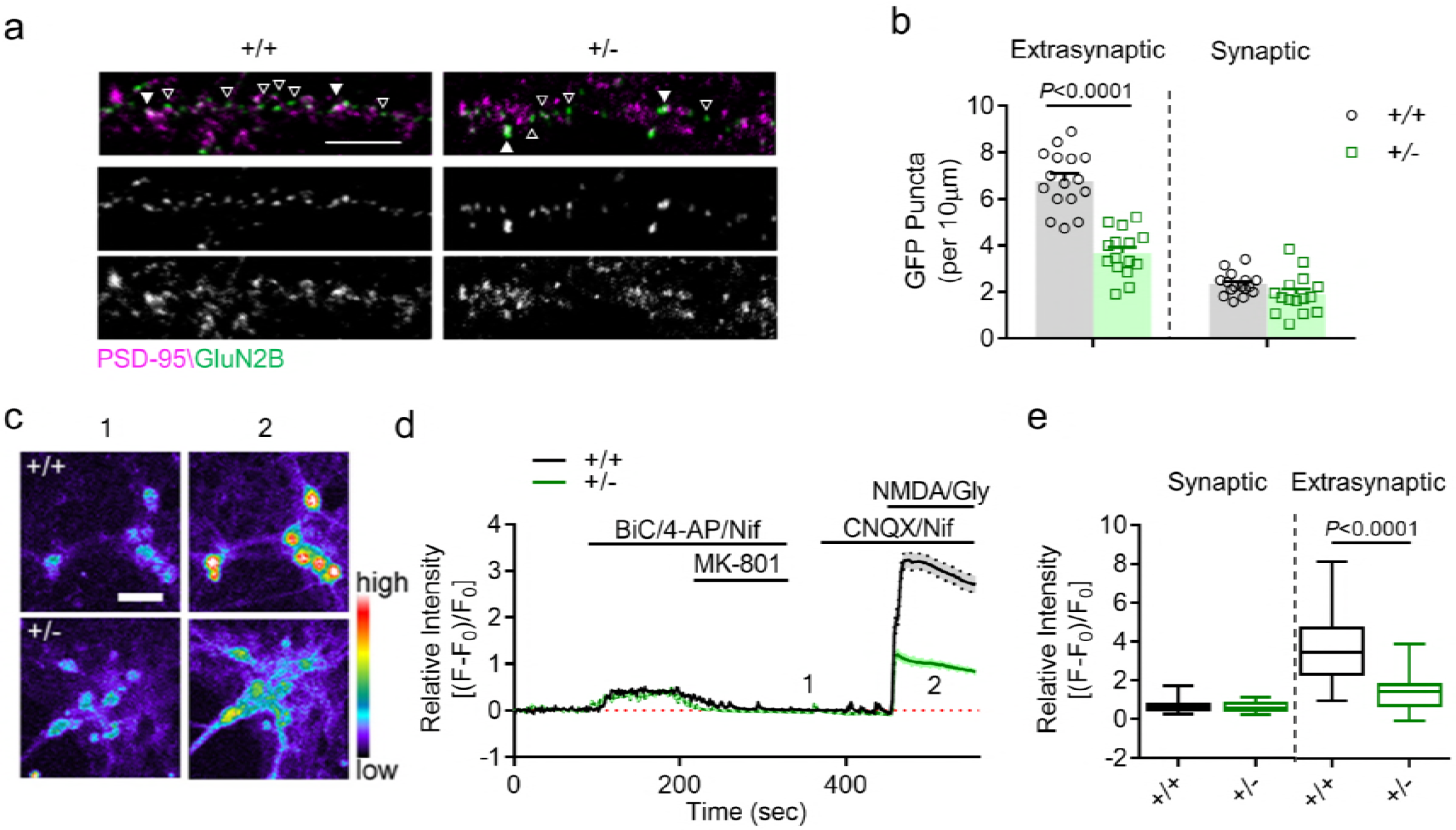
Kinesin-1 regulates extrasynaptic NMDAR surface targeting and function. (a) Representative images and (b) quantification of surface GFP-GluN2B co-localized with PSD-95 (n=16 for wild-type and n=15 for *kif5b*^+/−^ unpaired t-test; six images per genotype; scale bar = 10 μm). (c) Representative images, (d) time lapse, and (e) peak response (box plot) of calcium influx by extrasynaptic NMDA (exNMDA) receptor stimulation of *kif5b*^+/−^ (+/−) and wild-type (+/+) neurons. *Kif5b*^+/−^ (+/−) and wild-type (+/+) neurons evoked by Bic (20 μM)/4-AP (4 mM) in the presence of Nif (10 μM) and then blocked by MK-801 (10 μM), an irreversible NMDA receptor calcium channel blocker. The difference before and after MK-801 blockage was elicited by synaptic NMDARs. After 100 s of resting, NMDA and glycine were applied to activate the extrasynaptic NMDARs. The relative intensity (F-F_0_/F_0_) was calculated (n=65 for wild-type and n=64 for *kif5b*^+/−^ from two replicate experiments; e, Mann-Whitney U test; scale bar = 20 μm). Values are presented as mean ± s.e.m., unless otherwise indicated (test as indicated; exact *P* value given if less than 0.05).

### Kinesin-1 reduction attenuated NMDA-elicited excitotoxicity and ischemia-induced neurodegeneration

Nitric oxide (NO) is one of the main downstream molecules generated from NMDAR overactivation and acts as one of the main mediators for NMDAR induced excitotoxicity(Sattler et al., 1999). It is generated preferably from extrasynaptic NMDARs under certain neurodegenerative condition(Molokanova et al., 2014). We used an NO-sensitive electrode to measure NO emissions from acutely prepared hippocampal slices from wild-type and *kif5b*^+/−^ mice (Tjong, Jian et al., 2007). Hypoxia (N_2_) and L-Arginine induced significantly lower NO emissions in *kif5b*^+/−^ slices than in wild-type (L-Arg, *P*<0.0001, t= 9.53, df=9; N_2_, *P*<0.0001, t= 15.53, df=8; unpaired t-test; Supplementary Fig. S4a). Activation of NMDARs by NMDA (50 μM) or glutamate (Glu; 4 μM) together with glycine (Gly; 20 μM) increased NO emissions in both genotypes, but the amount was significantly less in *kif5b*^+/−^ slices (L-Glu, *P*<0.0001, t=27.10, df=17; NMDA, *P*<0.0001, t=29.26 df=6; unpaired t-test; Fig. S4, b and c). The NO emissions from both genotypes were greatly suppressed by NMDAR blocker MK-801 (100 μM) and NO synthase inhibitor L-NMMA (100 μM), indicating the NO emission was NMDAR-and NOS-dependent (Genotype x treatment, *P*<0.0001, F(2,19)=162.8; L-Glu vs. L-Glu/NMMA, *P*<0.0001, t=28.97, df=19; L-Glu vs. L-Glu/MK-801, *P*<0.0001, t=30.68, df=19; two-way ANOVA followed by Sidak *post hoc* test; Fig. S4d). Furthermore, in neurons with NMDARs blocked by MK-801 (10 μM), depolarization using KCl (50 mM) produced similar levels of NO emissions in both genotypes (KCl, *P*<0.0001, t=14.93, df=8; KCl/MK, *P*=0.1940, t=1.559, df=4; unpaired t-test; Fig. s4c). The reduced NO emission in *kif5b*^+/−^ slices might be caused by altered NO production machinery due to reduced kinesin-1, rather than reduced NMDAR-mediated calcium influx. To exclude this possibility, we first performed a Western blot analysis of nNOS in wild-type and *kif5b*^+/−^ slices, which showed identical amounts in both genotypes (*P*=0.3579, t=0.9689, df=9; unpaired t-test; Fig. S2c). Next, we treated wild-type and *kif5b*^+/−^ slices with A23187 (10 μM), which again showed similar NO emissions between genotypes (*P*=0.9155, t=0.1095, df=8; unpaired t-test; Fig. S4c).

As it is known that extrasynaptic NMDARs activation would consequently lead to exicitotoxicity, we tested whether the reduction of kinesin-1 would elicit neuroprotection against NMDA induced exicototoxicity. The heterozygous knockout of *kif5b* gene had minimal impact on the viability of the neuronal culture, as the neuronal morphology was indistinguishable from that of its wild-type littermates. We treated the cortical cultures from both wild-type and *kif5b*^+/−^ mice with NMDA (25 to 200 μM) and a co-agonist glycine (10 μM) for 10 minutes, which was more than sufficient to induce neuronal death. After NMDA washout, cortical neurons were cultured for an additional 24 hours. Cell death was examined by propidium iodide (PI) staining or by lactate dehydrogenase (LDH) assay, followed by morphological analysis under phase contrast microscopy. The *kif5b*^+/−^ neurons treated with 100 μM NMDA and 10 μM glycine exhibited significantly fewer PI-positive neurons and less membrane leakage compared with the wild-type group (*P*=0.0329, t=2.758, df=6; unpaired t-test; Fig. 4, a and b; NMDA *kif5b*^+/+^ vs. *kif5b*^+/−^, *P*<0.0001, t=8.575, df=21; two-way ANOVA followed by Sidak *post hoc* test; Fig. 4c). Morphologically, neurites in *kif5b*^+/−^ neurons were mostly intact indicating significantly less early neurodegeneration, whereas most neurons in the wild-type group showed fragmented neurites indicating synaptic loss leading to early neurodegeneration (*P*<0.0001, t=30.37, df=6; unpaired t-test; Fig. 4, a and b). Although NMDA elicited neurotoxicity in both genotypes in a dose-dependent manner, the neurotoxic effects were significantly lower in *kif5b*^+/−^ neurons at all tested concentrations (PI positive dose, F(2,17)=22.34, *P*<0.0001; degenerative dose, F(2,17)=10.63, *P*=0.0010; two-way ANOVA followed by Sidak *post hoc* test; Fig. S4, a and b). Moreover, treatment with memantine (50 μM), a potent NMDAR antagonist, significantly reduced NMDA-elicited neurotoxicity abolishing the differential neurotoxicity between *kif5b*^+/−^ and wild-type neurons (Genotype x Memantine, F(1,21)=38.64, *P*<0.0001; NMDA/Memantine +/+ vs. +/−, *P*=0.9797, t=0.7228, df=21; two-way ANOVA followed by Sidak *post hoc* test; Fig. 4c). Furthermore, selective inhibition of extrasynaptic NMDAR by a low dose of memantine (1 μM)(Okamoto et al., 2009) and GluN2B-specific inhibitor Ro25-6981 (5 μM) produced similar levels of NMDA-induced neurotoxicity in both wild-type and *kif5b*^+/−^ neurons (Mem, *P*=0.4627, t=0.7637, df=10; Ro25, *P*=0.7127, t=0.3789, df=10; unpaired t-test; Fig. S4d). As the control experiments, kainate (25 μM), a non-NMDA excitotoxin, produced similar neurotoxicity in *kif5b*^+/−^ and wild-type neurons (*P*=0.4883, t=0.7195, df=10; unpaired t-test; Fig. S4c). Treatment with A23187, a calcium ionophore that increases the calcium channel-independent level of intracellular calcium, elicited similar neurotoxicity in both genotypes (*P*=0.8807, t=0.1533, df=12; unpaired t-test; Fig. S4c), indicating kinesin-1 reduction elicited neuroprotection is selective to NMDAR and, at least in part, through preventing extrasynaptic NMDAR targeting and functioning.

**Figure 4.**
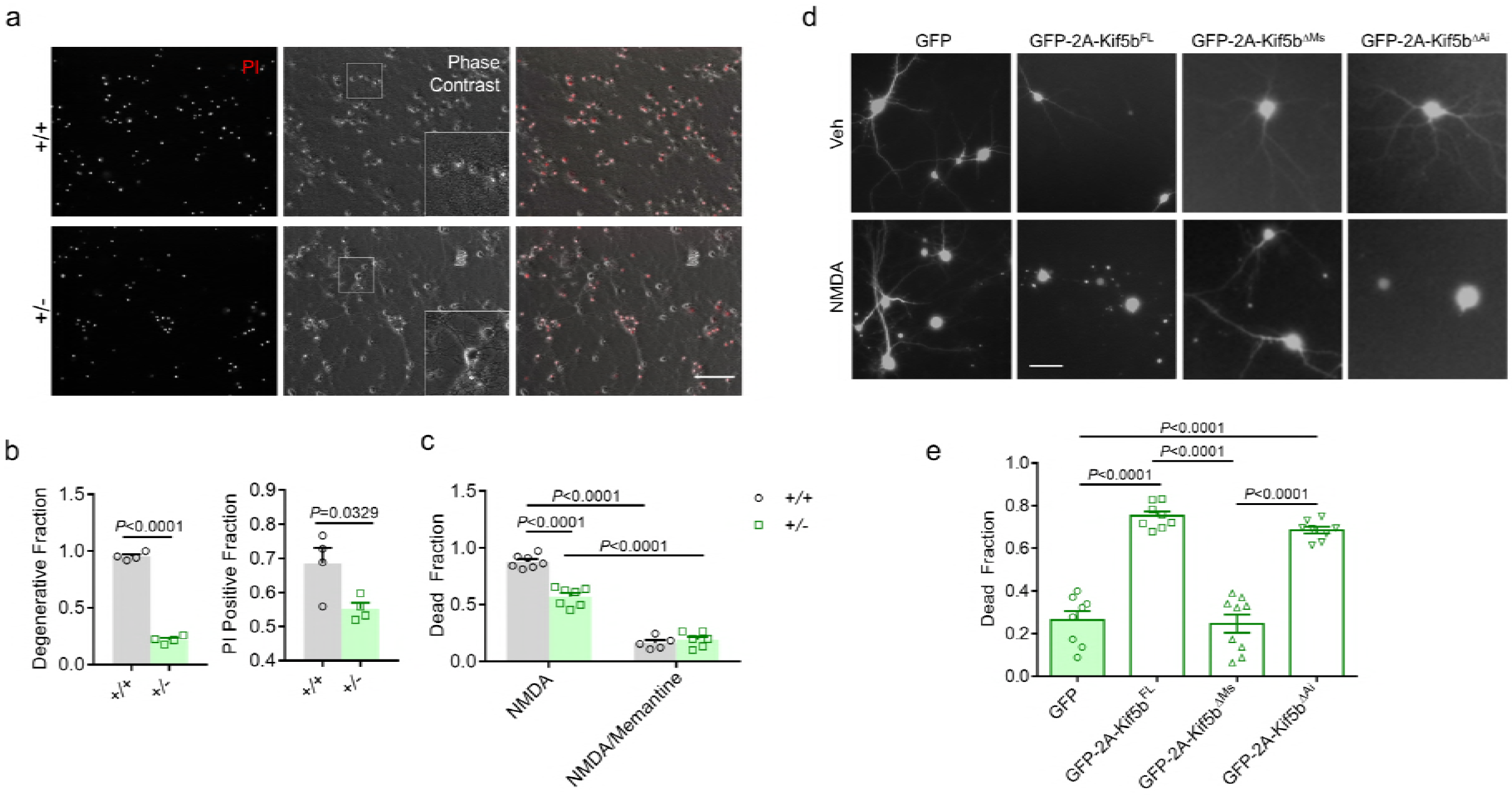
Kinesin-1 reduction protects neuron against NMDA induced exicitotoxicity. (a) Representative images and (b, left) quantification of PI-positive neurons and (b, right) degenerative neurons of wild-type (+/+) and *Kif5b*^+/−^ (+/−) cultures (n=4 per genotype, unpaired t-test) after treatment with NMDA/glycine (Gly) for 10 min followed by washing for 24 hours. The PI-positive fraction is the ratio of PI-positive neurons to PI-negative neurons, which indicates the portion of dead neurons. The degenerative fraction is the ratio of neurons with fragmented neurites to neurons with normal intact neurites, which indicates the portion of neurons with signs of early degeneration. Scale bar (white solid line) = 100 μm (50 μm in inset). (c) LDH leakage from NMDA-treated *kif5b*^+/−^ (+/−) and wild-type (+/+) neurons (n=7 for wild-type and n=6 for *kif5b*^+/−^ from three individual cultures, two-way ANOVA followed by Sidak *post hoc* test). The dead fraction is the ratio of LDH level in the drug-treated group to “all-kill” group. (d) Representative images and (e) quantification of dead and degenerative fractions of *kif5b*^+/−^ cultures transfected with GFP, GFP-2A-Kif5b^FL^, GFP-2A-Kif5b^ΔMT^, and GFP-2A-Kif5b^ΔAi^ after treatment with NMDA/Gly or vehicle (Veh) for 10 min followed by washing for 24 hours (n=8 for each genotype from three individual cultures, unpaired t-test). Scale bar (white solid line) = 50 μm. Values are presented as mean ± s.e.m. (test as indicated; exact *P* value given if less than 0.05).

To exclude the possibility that the reduced vulnerability to NMDA by reduced kinesin-1 level was due to secondary adaptation rather than the kinesin-1 reduction itself, we transiently overexpressed Kif5b with green fluorescent protein (GFP) separated by a 2A peptide in *kif5b*^+/−^ neurons. After NMDA treatment, *kifib*^+/−^ neurons transfected with GFP had well-preserved morphology with robust neurites, whereas neurons transfected with the full-length Kif5b-2A-GFP displayed shrunken nuclei with significant loss of neurites (*P*<0.0001, t=10.59, df=14; unpaired t-test; Fig. 4, d and e). Moreover, we assessed the NMDA-elicited cell death in *kif5b*^+/−^ neurons expressing either Kif5b with microtubule sliding site deletion or Kif5b with autoinhibitory site deletion. Transfection of either Kif5b mutants did not lead to spontaneous neurodegeneration (Fig. 3, d and e), as revealed by the intact neuronal morphology. We found only Kif5b with autoinhibitory site deletion had enhanced neuronal susceptibility to NMDA-elicited neurotoxicity, as shown by significantly more fragmented neurites (*P*<0.0001, t=9.079 df=15; unpaired t-test; Fig. 4, d and e). These results confirm the neuroprotection observed in *kif5b*^+/−^ neurons was due to a direct kinesin-1-mediated cellular mechanism, possibly associated with intracellular transportation.

To examine if the kinesin-1 reduction would affect neuronal fate following noxious insult *in vivo*, we performed BCCAO for 20 minutes on *kif5b*^+/−^ and wild-type mice. The *kif5b*^+/−^ mice were viable and fertile with no growth retardation (Genotype x time, male, F(5,18)=0.05272, *P*=0.9980, female, F(5,42)=1.088, *P*=0.3813; two-way ANOVA; Fig. S5a) and no signs of spontaneous neurodegeneration, as seen by the similar gross hippocampal anatomy in both genotypes (Fig. S5b). The induced transient global ischemia was sufficient to trigger neurodegeneration in the hippocampus (Tu et al., 2010). At 7 days after reperfusion, brain sections examined by TUNEL assay showed severe neuronal damage in the wild-type mice, but only mild to moderate damage in the *kif5b*^+/−^ mice. There was a significant increase in TUNEL-positive signals detected in wild-type hippocampal CA1 and CA3 regions (CA1, *P*<0.0001, t=6.0483, df=36; CA3, *P*<0.0001, t=10.7425, df=22; unpaired t-test; Fig. S5, c and d). Next, we performed MCAO for 120 minutes on *kif5b*^+/−^ and wild-type mice to induce focal cerebral ischemia (Lo, Chen et al., 2005, Tu et al., 2010). Consistent with BCCAO, we observed significantly improved neurological scores and reduced infarct volume in the *kifib*^+/−^ mice (Genotype, F (1, 11)=13.88, *P*=0.0033; two-way repeated measures ANOVA followed by Sidak *post hoc* test; infarct volume, *P*=0.0014, t=4.256, df=11; unpaired t-test; neurological score, U=6.000, *P*=0.0408, Mann-Whitney Rank Sum Test; Fig. 5, a to d).

**Figure 5.**
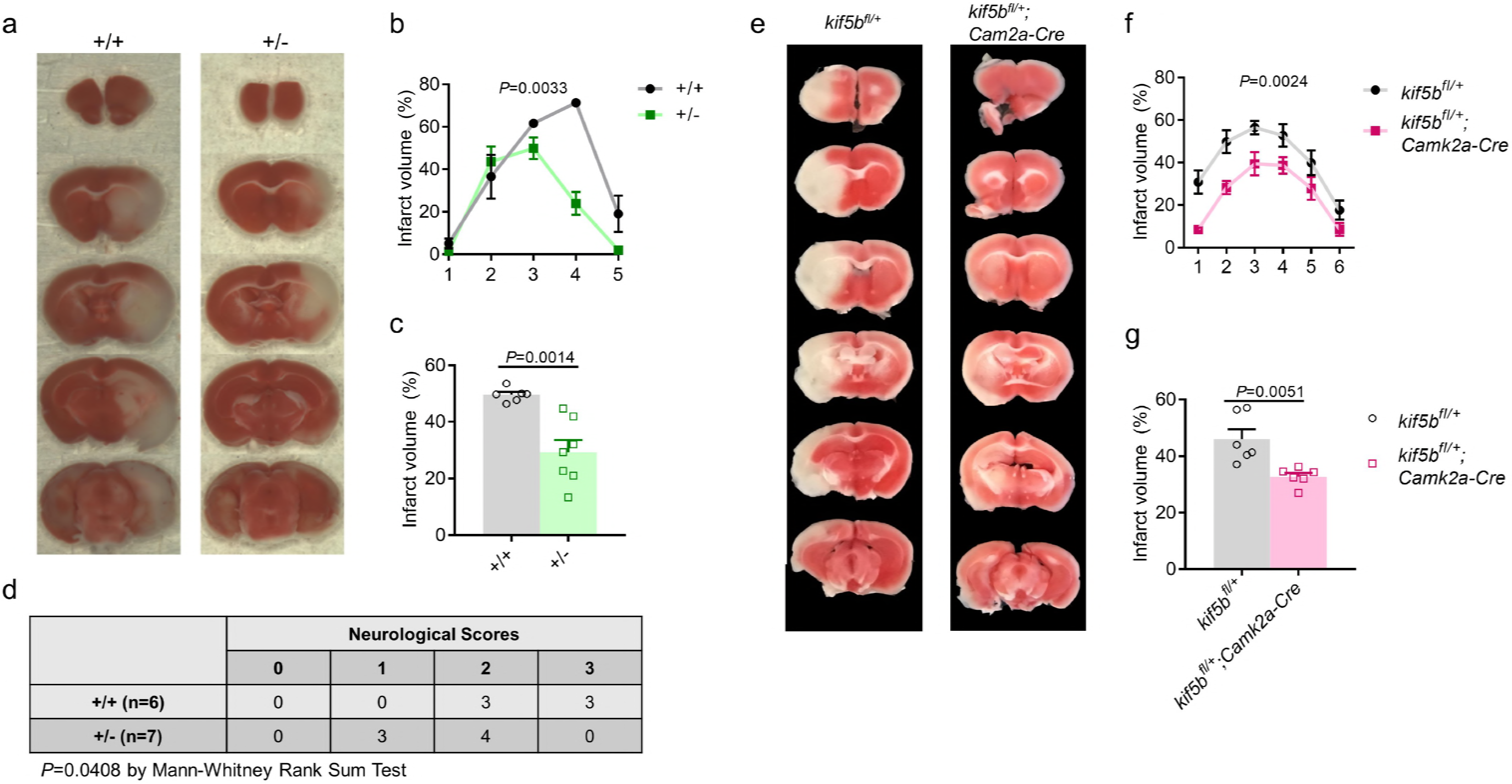
Kinesin-1 reduction protects brain against ischemia evoked neurodegeneration. (a) Representative images and (b and c) quantification of 2,3,5-triphenyltetrazolium chloride (TTC) staining of wild-type and *kif5b*^+/−^ mice after transient focal cerebral ischemia. Infarct volume of (b) the indicated slices and (c) whole brain (n=6 for wild-type and n=7 for *kif5b*^+/−^ mice; b, two-way ANOVA followed by Sidak *post hoc* test; c, unpaired t-test). (d) Neurological scoring revealed *kif5b*^+/−^ (+/−) mice had significant improvement in motor function after 2 hours ischemia and 24 hours reperfusion when compared to their wild-type littermates (+/+). (Neurological score: 0, freely moving; 1, reduced mobility; 2, partial paralysis; 3, no movement) (Mann-Whitney Rank Sum Test). (e) Representative images and (f and g) quantification of TTC staining of *kif5b*^*fl*/+^ and *kif5b*^*fl*/+^;*Camk2a-Cre* mice after MCAO. Infarct volume for indicated slices (f) and whole brain (g) (n=6 for *kif5b^fl/+^* and *kif5b*^*fl*/+^;*Camk2a-Cre*, respectively; f, two-way ANOVA followed by Sidak*post hoc* test; g, unpaired t-test). Values are presented as mean ± s.e.m., unless otherwise indicated (test as indicated; exact P value given if less than 0.05).

To examine the possibility that the observed neuroprotective phenotype in *kif5b*^+/−^ mice is directly due to kinesin-1 reduction rather than developmental adaptation *in vivo*, we generated a conditional knockout model by crossing *kif5b*^*fl*/+^ mice with Camk2a-Cre mice by *kif5b* excision starting on the third postnatal week to mimic postnatal Kif5b reduction, which should minimize developmental alteration caused by *kif5b* deletion (Tsien, Chen et al., 1996). As Camk2a-Cre mediates recombination only in a subset of excitatory neurons in the forebrain, it was not surprising to observe a modest reduction of Kif5b protein level at 6 weeks of age in both *kif5b*^fl/+^;*Camk2a-Cre* and *kif5b*^*fl/fl*^; *Camk2a-Cre* mice (F (2,6)=4.925, *P*=0.0543; oneway ANOVA; Fig. S5e). However, immunohistochemistry for Kif5b in the CA1 region, where Camk2a-Cre has the most impact, revealed a robust reduction in Kif5b protein level in *kif5b*^fl/+^;*Camk2a-Cre* mice when compared with their wide-type littermates (Fig. 5f). This data confirms that Kif5b was reduced at 6 weeks of age in a subset of neurons in the conditional knockout model. Consistent with our observation in *kifib*^+/−^ mice, the Camk2a-Cre-mediated conditional *kif5b* knockout mice showed robust resistance to MCAO-induced neurodegeneration (f, genotype, F (1, 10)=16.20, *P*=0.0024, two-way repeated measures ANOVA; g, *P*=0.0051, t=3.565 df=10; unpaired t-test; Fig. 5, e to g). Moreover, we observed a significant increase in NeuN immunoreactivity in the CA1 region of *kif5b* conditional knockout mice after 2 hours MCAO and 24 hours reperfusion when compared with their wild-type littermates (*P*=0.0022; Mann-Whitney Test; Fig. S5g). These combined results show that kinesin-1 reduction protects neurons against NMDAR-elicited excitotoxicity and ischemia-induced neurodegeneration.

### Reduction of Kif5b expression associated with suppression of cell death pathways

Ischemia precondition (IPC) elicited neural activity dependent neuroprotection by altering the transcriptome to adapt and resist subsequent noxious insult(Stenzel-Poore, Stevens et al., 2003). As previous findings show a robust neuroprotective effect by kinesin-1 reduction, we hypothesized that the alteration of kinesin genes, particularly kinesin-1, might be a core mechanism underlying the IPC elicited neuroprotection. To test this, we analyzed a microarray dataset (GSE32529) that profiled gene expressions at 3, 24 and 72 hours after a brief cerebral ischemia to induce ischemia preconditioning (IPC) in mice. Significantly altered genes at 3, 24, and 72 hours after IPC were filtered and analyzed by Ingenuity Pathway Analysis (IPA) platform (Qiagen) and compared with sham control groups (Fig. 6a). By combining and comparing results from the “Diseases and Functions” analysis at the three indicated time points, we found a set of genes relating to apoptosis and cell death were robustly enriched at 3 hours after IPC, while this set of genes was suppressed at 24 and 72 hours after IPC (Fig. 6b). The finding of enhanced genes relating to apoptosis and cell death at 3 hours after IPC was unexpected, and the overall trend supports the notion that brief exposure to IPC reprograms gene expression to protect the brain against lateral noxious insult. Besides the expected trends of inhibited cell death pathways/genes at the extended times after IPC, we unexpectedly identified suppressed genes related to neurodevelopment at over 3 to 72 hours (Fig. 6b). This raises the possibility that the same set of genes that regulate neurodevelopment also play a key role in mediating neuronal adaptation to stress, including noxious insult. Kinesin genes plays a critical role in the development of cells and tissues such as neurons, muscles, chondrocytes, and osteoblasts (Hsu, Zhang et al., 2011, Muhia, Thies et al., 2016, Padamsey, McGuinness et al., 2017, Qiu, Xiao et al., 2012, Wang, Cui et al., 2013). We therefore investigated the global expression profile of kinesin genes at 3, 24, and 72 hours after IPC in mice compared to the sham control group. At 3 hours after IPC, the cumulative distribution of kinesin genes in the differentially expressed population exhibited a robust shift toward upregulation (D=0.2779, *P*=0.0009, Kolmogorov-Smirnov test; Fig. S6a), while this trend disappeared after 24 hours (D=0.1387, *P*=0.0891; Kolmogorov-Smirnov test; Fig. S6a). In contrast, the cumulative distribution of kinesin genes exhibited a significant shift toward downregulation at 72 hours compared with the overall gene expression profile (D= 0.1583, *P*= 0.0347; Kolmogorov-Smirnov test; Fig. 6c). This became more apparent when comparing the cumulative expression profile of kinesin genes at 3 hours and at 72 hours, which showed a significant downregulation shift (D= 0.2785, *P*=0.0165, Kolmogorov-Smirnov test; Fig. 6d). The results of the cumulative expression profile of kinesin genes suggests a time-dependent downregulation shift after IPC, which might indicate a kinesin-dependent mechanism underlying IPC-induced adaptation and neuroprotection. To identify which kinesin is the key player, we extracted probes for kinesin genes that were robustly altered at 72 hours after IPC, which revealed two probes for *kif5b*. Moreover, we found *kif5b* and *kif1b* expressions, but not *kif3a*, were robustly repressed at 72 hours compared to at 3 hours after IPC (*kif5b*, *P*= 0.0313; *kif1b*, *P*=0.0313; *kif3a*, *P*=0.1250; Wilcoxon matched-pairs signed rank test; Fig. S6b), suggesting repression of *Kif5b* gene was a candidate event regulating neuronal adaptation to further ischemia insult.

**Figure 6.**
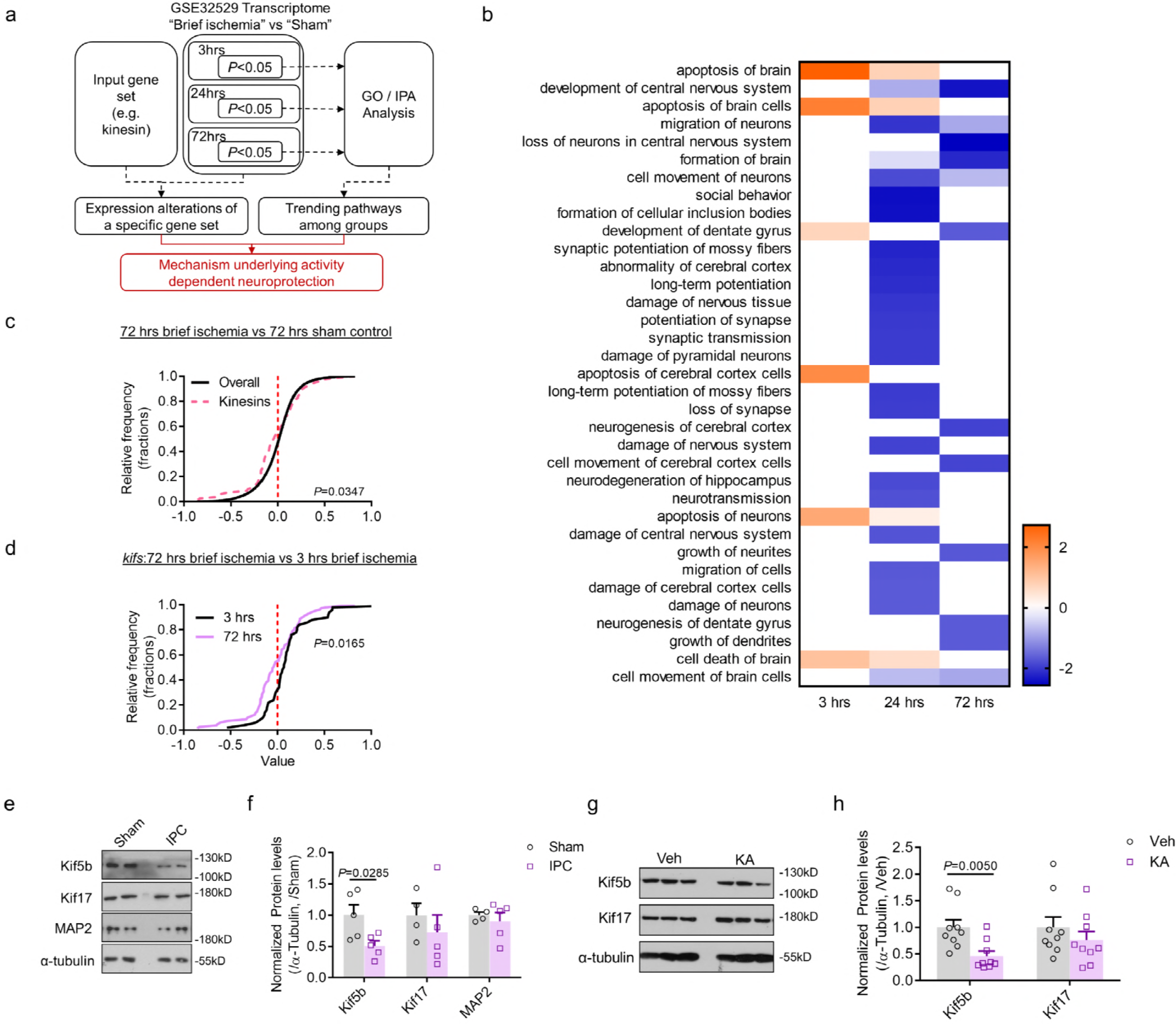
Transcriptome analysis of mouse cortex after ischemia preconditioning reveals the suppression of cell death pathways is associated with reduced *kif5b* expression. (a) Schematic figure of the transcriptome analysis. The genes significantly altered at 3, 24, or 72 hours (hrs) after brief cerebral ischemia preconditioning (IPC) were analyzed by Ingenuity Pathway Analysis (IPA) platform. The expression of kinesin genes (*kifs*) were extracted and compared with the overall expression profile. (b) IPA results of the pathways categorized by diseases and functions. (c) The expression profiling of kinesin genes (*kifs*) were compared with the overall gene expression profile at 72 hrs after IPC normalized to 72 hrs in the sham control (Kolmogorov-Smirnov test). (d) The expression profiling of kinesin genes at 3 and 72 hrs after IPC (Kolmogorov-Smirnov test). (e) Representative Western blot images and (f) quantification of Kif5b, Kif17, and MAP2 in the hippocampus of wild-type mice after IPC or sham operation (n=5 for the IPC group and n=4 for the sham operation group, unpaired t-test). (g) Representative Western blot image and (h) quantification of Kif5b, Kif17, and MAP2 in the hippocampus of wild-type mice with kainate treatment (KA, 12 mg/kg, 2 hrs) or vehicle (n=9 for the KA group and n=9 for the vehicle group, unpaired t-test). Values are presented as mean ± s.e.m. (test as indicated; exact *P* value given if less than 0.05).

To validate the transcriptome analysis, we first examined the expression levels of transport-related genes in a mouse IPC model. At 48 hours after reperfusion, the level of Kif5b, the heavy chain of the microtubule-dependent motor kinesin-1, was significantly reduced (*P*=0.0069, t=3.605, df=8, unpaired t-test). However, levels of Kif17 and microtubule-associated protein-2 (MAP2), a component of neuron microtubule, were not changed (Kif17, *P*=0.4703, t=0.7632, df=7; MAP2, *P*=0.5731, t=0.5910, df=7; unpaired t-test; Fig. 6, e and f). Next, we tested whether downregulation of Kif5b was specific to IPC or whether this was a response to the global regulation of neuronal activity. Mice injected with kainate, which acts as an inhibitor of synaptic GABA, resulted in increased neuronal activity (Ramamoorthi, Fropf et al., 2011). We found the level of Kif5b, but not Kif17, was reduced at 48 hours after kainate treatment (Kif5b, *P*=0.005, t=3.2533, df=16; Kif17, *P*=0.3552, t=0.9522, df=16; unpaired t-test; Fig. 6, g and h), which indicates the reduction of kinesin-1 was associated with increased neuronal activity. This suggests a role of the intracellular transportation system underlying the suppression of cell death pathways by IPC and indicates that kinesin-1 dependent alteration of NMDAR compartmentalization might be the core mechanism.

## Discussion

The present study revealed novel complex between kinesin-1 and GluN2B-containing NMDARs. This complex mediates the extrasynaptic targeting of NMDARs, the resulting pro-death signaling including calcium influx and NO production, and eventually mediates neuronal vulnerability to NMDA-induced excitotoxicity and ischemia-evoked neurodegeneration. The transcriptome of IPC further implicates a role of kinesin-1 in mediating activity dependent neuroprotection, probably via altering NMDAR compartmentalization and shifting the balance to highlight pro-survival pathways. Our findings demonstrate a role of Kinesin-1 in mediating the compartmentalization of signaling pathways within neurons, which can be tuned down by suppressing Kinesin-1 expression or by dissociating the interaction between Kinesin-1 and GluN2B containing NMDAR when neurons sense a detrimental insult.

It has long been known that hyperactivation of NMDARs contribute to excitotoxicity and neuronal death. NMDAR-related excitotoxicity also acts as the major mechanism that causes neuronal death in both acute and chronic neurodegenerative disorders. However, attempts to translate NMDAR antagonization to clinic applications are far from satisfactory(De Keyser et al., 1999). The reasons might stem from the complexity of NMDAR-mediated signaling and subcellular compartmentalization(Zhou & Sheng, 2013). There are two distinct pools of NMDAR: synaptic NMDAR steadily accumulates at the postsynaptic density, whereas extrasynaptic NMDAR is more mobile and dynamic in extrasynaptic regions. The downstream signaling and the functional consequence of these two distinct pools of NMDAR are also different. The activation of synaptic NMDAR activates MAPK signaling and modulates synaptic strength(Thomas & Huganir, 2004). The dysfunction of this pathway not only causes learning and memory deficits, but also neuropsychiatric disorders such as schizophrenia(Cohen, Tsien et al.). The activation of synaptic NMDAR is reported to antagonize the calcium-elicited pro-death pathway(Hardingham et al., 2002). Our knowledge of extrasynaptic NMDAR is far less thorough compared to our understanding of synaptic NMDAR. The activation of extrasynaptic NMDAR was found to suppress the pro-death pathway mediated by synaptic NMDAR, which shifts the balance toward cell death in neurons. Furthermore, DARK1 is recruited to GluN2B and mediates the pro-death signaling toward excitotoxicity(Tu et al., 2010), while PGC-1α opposes extrasynaptic NMDARs function(Puddifoot, Martel et al., 2012).

One major question in this field is how the compartmentalization of signaling is regulated. Previous evidence indicates that extrasynaptic NMDARs are more mobile than the synaptic NMDARs(Groc, Heine et al., 2004), which makes the convergence of different transportation systems possible. In support of this notion, dynamin-dependent regulation is known to limit extrasynaptic synaptic activation(Wild, Jones et al., 2014a). Our finding that microtubule-dependent transportation relies on extrasynaptic NMDAR fits into current model in two ways. First, Kinesin-1 may directly convey GluN2B-containing NMDAR to the extrasynaptic region. Kinesin-1 has been known to directly interact with Myosin Va(Huang, Brady et al., 1999), suggesting cooperation between microtubule-dependent transportation and actin-dependent transportation. Besides synaptic regions, actin filament dynamics are found to be enriched at the lateral zone and neck of the spine(Frost, Shroff et al.). Our model suggests that extrasynaptic NMDAR is first conveyed and targeted to extrasynaptic regions by microtubule-dependent transport, and then moves between synaptic and extrasynaptic regions through the cooperation of microtubule-dependent and actin-dependent transport systems. Second, Kinesin-1 may convey internalized NMDAR from synaptic regions to extrasynaptic regions(Du, Su et al., 2016). This is supported by the evidence that internalized NMDARs contain vesicles that are attached to microtubule-like filaments(Tovar & Westbrook, 1999), which further indicates a role of Kinesin-1 in determining the fate of the internalized vesicles towards recycling or degradation.

Our finding that Kinesin-1 mediates transport of NMDAR adds complexity to the anterograde transport of NMDAR. It has been proposed that Kif17 drives NR2B-containing NMDA receptors to synaptic sites(Setou et al., 2000, Yin et al., 2011), which is in parallel with our observation that knockdown of Kif5b reduced only extrasynaptic targeting and functioning. It is widely recognized that neurons may use different transport systems to convey cargo to distinct sub-domains, at least in the case of mitochondria transport, which relies on both Kif5 and Kif1b(Nangaku, Sato-Yoshitake et al., 1994, Wang & Schwarz, 2009). When reaching the site with high levels of calcium, the mitochondrial membrane protein Miro mediates detachment or folding of kinesin-1, which traps the mitochondria(Wang & Schwarz, 2009), whereas Kif1b uses a KBP-dependent mechanism that is distinct from kinesin-1(Wozniak, Melzer et al., 2005). NMDAR exists as both diheteromer consisting of GluN1/GluN2A or GluN1/GluN2B as the majority, or triheteromer consisting of GluN1/GluN2A/GluN2B *in vivo*. The functional property of these two forms of NMDAR are different(Hansen, Ogden et al., 2014). The triheteromer is presumably dominant form of NMDAR in the adult forebrain, consisting with our data showing that kinesin-1 co-immunoprecipitates with both GluN2B and GluN2A (Fig. 1a). However, it has also been reported that GluN2B containing NMDAR is the major form of NMDAR in the extrasynaptic site(Tovar & Westbrook, 1999). It is plausible that kinesin-1 has greater affinity to the diheteromer consisting of GluN1/GluN2B, which therefore enriches GluN2B-containing NMDAR in the extrasynaptic region, while the Kif17 has greater impact on triheteromer, which is enriched in the synaptic region. Supporting this notion, our analysis on the surface distribution NMDAR reveals the reduction of GluN2B, but not GluN2A, in the surface (Fig. 1, g and h). To understand the function and compartmentalization of NMDARs in more detail, especially its role in activity-dependent plasticity and in disease pathology, we need to examine the regulatory or coordinating mechanisms among the different kinesins and between different transportation systems.

It is well established that the hierarchical assembly of synapses and neuronal contacts is highly dependent on intracellular transportation systems(Hirokawa et al., 2010). Our findings provide a novel intrinsic activity-dependent mechanism for the regulation of NMDARs, which protects neurons against excitotoxic insult. Reduction of NMDAR activity could be regarded as a form of protective feedback that limits excess NMDAR-dependent calcium influx triggering excitotoxicity. This feedback can be through a number of factors, such as NMDAR subunit composition(Martel, Ryan et al., 2012), the association of scaffolding proteins(Forder & Tymianski, 2009), and neuronal activity(Groc et al., 2004, Wild, Jones et al., 2014b). Furthermore, our findings combine and unify the associations between kinesin level, elicited neuroprotection, and subsequent impact on extrasynaptic NMDAR, which support the notion that regulating components beyond the synapse might be essential to induce neuroprotection. Consistent with our current findings, reduced mTORC1 activity and the resulting induction of autophagy has been proposed as another neuronal activity-dependent mechanism to induce tolerance. Interestingly, this mechanism might function closely with intracellular transportation systems in eliciting neuroprotection(Fu, Nirschl et al., 2014, Papadakis, Hadley et al., 2013). Altogether, these findings provide a potential therapeutic strategy that could delay degenerative processes by extending the period and effect of intrinsic neuroprotective mechanisms.

Inhibiting kinesin has been proposed as a promising approach to treating cancer, and some small molecules have already been developed. For example, the Kif11 motor protein mediates mitosis and drives glioblastoma invasion, proliferation, and self-renewal. Inhibition of Kif11 by a specific small molecule could stop the progression of glioblastoma(Venere, Horbinski et al., 2015). The perturbation of intracellular transport may not only treat malignant dividing cells such as cancer, but also show promising effects on the central nervous system (CNS). Epothilone D, a microtubule stabilizer that can reduce microtubule dynamic and dependent transport, was able to ameliorate the impaired exploratory behavior in a Rett mouse model(Delépine, Meziane et al., 2016). Epothilone B, an FDA approved microtubule stabilizer that can limit microtubule-dependent transport, was able to reactivate neuronal polarization and promote axon regeneration after CNS injury(Ruschel, Hellal et al., 2015).

Interestingly, *in-situ* detection of *kif5b* mRNA reveals high expressions of Kif5b in the hippocampus, cortex, pons and medulla, but not striatum, hypothalamus or midbrain(LeinHawrylycz et al., 2006, Thomas & Huganir, 2004). Extrasynaptic NMDAR has been implicated in both acute and chronic neurodegenerative diseases such as stroke and Alzheimer’s disease. This makes Kif5b a potential therapeutic candidate, because Kif5b inhibition would have a robust impact on regions including the hippocampus where neurodegeneration mostly occurs, but a minimal impact on other critical brain regions including the striatum that are less affected by excitotoxicity. Moreover, suppressing Kif5b expression, particularly by disrupting the Kif5b-GluN2B interaction, might open new therapeutic avenues. As the structure for kinesin-1 cargo binding motif has been resolved(Pernigo, Lamprecht et al., 2013, Yip, Pernigo et al., 2016), it would be tempting to resolve the structure for kinesin-1-GluN2B complex and use this structure as the basis to predict or design molecules that could halt extrasynaptic NMDAR mediated neurodegeneration. As such an approach would mimic intrinsic changes induced by brief and beneficial stimulations, it might be tolerated by brain and thus minimize potential side effects. This would also prolong the period before the brain becomes vulnerable to detrimental insult, enabling the brain to adapt or develop resistance(Stenzel-Poore et al., 2003), delay functional impacts like memory or motor deficits, and allow for efficient interventions(Armstead, Hekierski et al., 2018).

These reports suggest targeting of intracellular transport, including kinesin-1, could be potential targets for deferring neurodegeneration. Compared with previous approaches of NMDAR antagonization, modulating kinesin-1-dependent transport could extend or reproduce native pro-life signaling, which could effectively hinder neurodegeneration while minimizing side effects. Thus, we propose that the fine-tuning of intracellular transportation could be a promising translational target for neurodegenerative disorders.

## Materials and Methods

All experimental protocols were reviewed and approved by the Committee on the Use of Live Animals in Teaching and Research, the University of Hong Kong. The Kif5b^+/−^ mice were generated by standard gene targeting via homologous recombination in mouse embryonic stem cells (Cui, Wang et al., 2011). Please refer to the Materials and Methods section in the supplementary materials for more details.

## Author contributions

The work presented here was carried out in collaboration with all authors. J.D.H. directed and coordinated the study. R.L. designed and conducted the experiments and analyzed the data. Z.D. performed the GST pull-down experiments and rescue experiments. J.W. performed the pilot experiments. M.L.F., C.F.L., and Y.L. conducted the NO measurements. D.Y., A.C.Y.L., H.C., H.S., and J. S. helped with the MCAO experiments. Y.N., Z.W., and J.C. helped with the GST pull-down experiments and animal maintenance. W.W., J.X., W.H.Y., and Y.S.C. provided critical comments on the project. R.L. and J.D.H. wrote the manuscript. All authors read, commented on, and approved the manuscript.

The authors have declared that no conflict of interest exists.

## Acknowledgments

We thank Dr. Jianhong Luo for providing the GFP-GluN2B plasmid, Dr. Anne Stephenson for providing the GluN1-1a plasmid, and Dr. Scott Brady and Dr. Gerardo Morfini for providing the mouse Kif5b cDNA. We thank Dr. Jing Guo, Mr. Cyril Lai, and Miss Jess Chan for their technical support for the confocal microscopy. We thank Dr. Yue Zhuo (Shenzhen Institutes of Advanced Technology, Chinese Academy of Sciences) for helping in the characterization of protein-protein interactions. This study was supported by the University Grants Committee of Hong Kong (AoE/M-04/04), Hong Kong Research Grants Council (HKU 767012/04M, HKU 768113M, HKU 17127015/05M, HKUST10/CRF/12R), Hong Kong Scholars Program (No. XJ2016055), National Basic Research Program of China (973 Program 2014CB745200) of the Ministry of Science and Technology of PRC, Shenzhen Peacock project (KQTD2015033117210153), and Shenzhen Science and Technology Innovation Committee Basic Science Research Grant (JCYJ20150629151046896).

## The paper explained

### PROBLEM

Neurodegenerative disorders, chronic or acute, are prevailing and are putting mounting pressure on health system. The hyperactivation of NMDARs is regarded as the common mechanism contributing to cell death among different neurodegenerative disorders, however, the translation of NMDAR inhibition to halt neurodegeneration is far from satisfaction.

### RESULT

We found that Kif5b, the heavy chain of a microtubule dependent motor kinesin-1, directly bound with GluN2B. This binding regulated the targeting and function of extrasynaptic NMDAR, a specific pool of NMDAR mediating cell death. More importantly, partial reduction of Kif5b reduced kinesin-1-NMDAR complex level *in vivo*, protected neurons against NMDA-elicited excitotoxicity, and protected brain against ischemia-evoked neurodegeneration. The reduction of kinesin dependent transport was an activity dependent resilient mechanism as we found a global reduction of kinesin motors, including kinesin-1, in the cerebral ischemia preconditioning transcriptome.

### IMPACT

Our finding reveals a molecular complex specific regulates the pro-death pool of NMDARs, and the reduction of kinesin-1 is an intrinsic neuroprotective mechanism and is tolerated *in vivo*. It is indicative that manipulation of kinesin-1-NMDAR interaction could be exploited to halt the progression of neurodegeneration that is caused by NMDAR hyperactivation, which could circumvent the potential side effects caused by conventional and global NMDAR inhibition.

## References

Armstead WM, Hekierski H, Pastor P, Yarovoi S, Higazi AA-R, Cines DB (2018) Release of IL-6 After Stroke Contributes to Impaired Cerebral Autoregulation and Hippocampal Neuronal Necrosis Through NMDA Receptor Activation and Upregulation of ET-1 and JNK. Translational stroke research: 1–8

Bading H (2013) Nuclear calcium signalling in the regulation of brain function. Nat Rev Neurosci 14: 593–608

Bading H (2017) Therapeutic targeting of the pathological triad of extrasynaptic NMDA receptor signaling in neurodegenerations. The Journal of Experimental Medicine 214: 569–578

Chu Y, Morfini GA, Langhamer LB, He Y, Brady ST, Kordower JH (2012) Alterations in axonal transport motor proteins in sporadic and experimental Parkinson’s disease. Brain 135: 2058–2073

Cohen SM, Tsien RW, Goff DC, Halassa MM The impact of NMDA receptor hypofunction on GABAergic neurons in the pathophysiology of schizophrenia. Schizophrenia Research 167: 98–107

Cui J, Wang Z, Cheng Q, Lin R, Zhang XM, Leung PS, Copeland NG, Jenkins NA, Yao KM, Huang JD (2011) Targeted inactivation of kinesin-1 in pancreatic beta-cells in vivo leads to insulin secretory deficiency. Diabetes 60: 320–30

De Keyser J, Sulter G, Luiten PG (1999) Clinical trials with neuroprotective drugs in acute ischaemic stroke: are we doing the right thing? Trends Neurosci 22: 535–40

Delépine C, Meziane H, Nectoux J, Opitz M, Smith AB, Ballatore C, Saillour Y, Bennaceur-Griscelli A, Chang Q, Williams EC, Dahan M, Duboin A, Billuart P, Herault Y, Bienvenu T (2016) Altered microtubule dynamics and vesicular transport in mouse and human MeCP2-deficient astrocytes. Human Molecular Genetics 25: 146–157

Du W, Su QP, Chen Y, Zhu Y, Jiang D, Rong Y, Zhang S, Zhang Y, Ren H, Zhang C, Wang X, Gao N, Wang Y, Sun L, Sun Y, Yu L (2016) Kinesin 1 Drives Autolysosome Tubulation. Dev Cell 37: 326–36

Forder JP, Tymianski M (2009) Postsynaptic mechanisms of excitotoxicity: Involvement of postsynaptic density proteins, radicals, and oxidant molecules. Neuroscience 158: 293–300

Frost NA, Shroff H, Kong H, Betzig E, Blanpied TA Single-Molecule Discrimination of Discrete Perisynaptic and Distributed Sites of Actin Filament Assembly within Dendritic Spines. Neuron 67: 86–99

Fu MM, Nirschl JJ, Holzbaur EL (2014) LC3 binding to the scaffolding protein JIP1 regulates processive dynein-driven transport of autophagosomes. Dev Cell 29: 577–90

Groc L, Heine M, Cognet L, Brickley K, Stephenson FA, Lounis B, Choquet D (2004) Differential activity-dependent regulation of the lateral mobilities of AMPA and NMDA receptors. Nat Neurosci 7: 695–6

Hansen Kasper B, Ogden Kevin K, Yuan H, Traynelis Stephen F (2014) Distinct Functional and Pharmacological Properties of Triheteromeric GluN1/GluN2A/GluN2B NMDA Receptors. Neuron 81: 1084–1096

Hardingham GE, Fukunaga Y, Bading H (2002) Extrasynaptic NMDARs oppose synaptic NMDARs by triggering CREB shut-off and cell death pathways. Nat Neurosci 5: 405–14

Hirokawa N, Niwa S, Tanaka Y (2010) Molecular Motors in Neurons: Transport Mechanisms and Roles in Brain Function, Development, and Disease. Neuron 68: 610–638

Hsu S-HC, Zhang X, Yu C, Li ZJ, Wunder JS, Hui C-C, Alman BA (2011) Kif7 promotes hedgehog signaling in growth plate chondrocytes by restricting the inhibitory function of Sufu. Development 138: 3791–3801

Huang J-D, Brady ST, Richards BW, Stenoien D, Resau JH, Copeland NG, Jenkins NA (1999) Direct interaction of microtubule-and actin-based transport motors. Nature 397: 267

Kaan HY, Hackney DD, Kozielski F (2011) The structure of the kinesin-1 motor-tail complex reveals the mechanism of autoinhibition. Science 333: 883–5

Lein ES, Hawrylycz MJ, Ao N, Ayres M, Bensinger A, Bernard A, Boe AF, Boguski MS, Brockway KS, Byrnes EJ, Chen L, Chen L, Chen T-M, Chi Chin M, Chong J, Crook BE, Czaplinska A, Dang CN, Datta S, Dee NR et al. (2006) Genome-wide atlas of gene expression in the adult mouse brain. Nature 445: 168

Lo AC, Chen AY, Hung VK, Yaw LP, Fung MK, Ho MC, Tsang MC, Chung SS, Chung SK (2005) Endothelin-1 overexpression leads to further water accumulation and brain edema after middle cerebral artery occlusion via aquaporin 4 expression in astrocytic end-feet. J Cereb Blood Flow Metab 25: 998–1011

Martel MA, Ryan TJ, Bell KF, Fowler JH, McMahon A, Al-Mubarak B, Komiyama NH, Horsburgh K, Kind PC, Grant SG, Wyllie DJ, Hardingham GE (2012) The subtype of GluN2 C-terminal domain determines the response to excitotoxic insults. Neuron 74: 543–56

Molokanova E, Akhtar MW, Sanz-Blasco S, Tu S, Piña-Crespo JC, McKercher SR, Lipton SA (2014) Differential Effects of Synaptic and Extrasynaptic NMDA Receptors on Aβ-Induced Nitric Oxide Production in Cerebrocortical Neurons. The Journal of Neuroscience 34: 5023–5028

Muhia M, Thies E, Labonté D, Ghiretti AE, Gromova KV, Xompero F, Lappe-Siefke C, Hermans-Borgmeyer I, Kuhl D, Schweizer M, Ohana O, Schwarz JR, Holzbaur ELF, Kneussel M (2016) The Kinesin KIF21B Regulates Microtubule Dynamics and Is Essential for Neuronal Morphology, Synapse Function, and Learning and Memory. Cell Reports 15: 968–977

Murrough JW, Abdallah CG, Mathew SJ (2017) Targeting glutamate signalling in depression: progress and prospects. Nature Reviews Drug Discovery 16: 472

Nangaku M, Sato-Yoshitake R, Okada Y, Noda Y, Takemura R, Yamazaki H, Hirokawa N (1994) KIF1B, a novel microtubule plus end-directed monomeric motor protein for transport of mitochondria. Cell 79: 1209–1220

Okamoto S, Pouladi MA, Talantova M, Yao D, Xia P, Ehrnhoefer DE, Zaidi R, Clemente A, Kaul M, Graham RK, Zhang D, Vincent Chen HS, Tong G, Hayden MR, Lipton SA (2009) Balance between synaptic versus extrasynaptic NMDA receptor activity influences inclusions and neurotoxicity of mutant huntingtin. Nat Med 15: 1407–13

Padamsey Z, McGuinness L, Bardo SJ, Reinhart M, Tong R, Hedegaard A, Hart ML, Emptage NJ (2017) Activity-Dependent Exocytosis of Lysosomes Regulates the Structural Plasticity of Dendritic Spines. Neuron 93: 132–146

Paoletti P, Bellone C, Zhou Q (2013) NMDA receptor subunit diversity: impact on receptor properties, synaptic plasticity and disease. Nature Reviews Neuroscience 14: 383

Papadakis M, Hadley G, Xilouri M, Hoyte LC, Nagel S, McMenamin MM, Tsaknakis G, Watt SM, Drakesmith CW, Chen R, Wood MJ, Zhao Z, Kessler B, Vekrellis K, Buchan AM (2013) Tsc1 (hamartin) confers neuroprotection against ischemia by inducing autophagy. Nat Med 19: 351–7

Pernigo S, Lamprecht A, Steiner RA, Dodding MP (2013) Structural Basis for Kinesin-1:Cargo Recognition. Science 340: 356–359

Puddifoot C, Martel M-A, Soriano FX, Camacho A, Vidal-Puig A, Wyllie DJA, Hardingham GE (2012) PGC-1α Negatively Regulates Extrasynaptic NMDAR Activity and Excitotoxicity. The Journal of Neuroscience 32: 6995–7000

Qiu N, Xiao Z, Cao L, Buechel MM, David V, Roan E, Quarles LD (2012) Disruption of Kif3a in osteoblasts results in defective bone formation and osteopenia. Journal of Cell Science 125: 1945–1957

Ramamoorthi K, Fropf R, Belfort GM, Fitzmaurice HL, McKinney RM, Neve RL, Otto T, Lin Y (2011) Npas4 regulates a transcriptional program in CA3 required for contextual memory formation. Science 334: 1669–75

Ruschel J, Hellal F, Flynn KC, Dupraz S, Elliott DA, Tedeschi A, Bates M, Sliwinski C, Brook G, Dobrindt K, Peitz M, Brüstle O, Norenberg MD, Blesch A, Weidner N, Bunge MB, Bixby JL, Bradke F (2015) Systemic administration of epothilone B promotes axon regeneration after spinal cord injury. Science 348: 347–352

Sanacora G, Frye MA, McDonald W, et al. (2017) A consensus statement on the use of ketamine in the treatment of mood disorders. JAMA Psychiatry 74: 399–405

Sattler R, Xiong Z, Lu WY, Hafner M, MacDonald JF, Tymianski M (1999) Specific coupling of NMDA receptor activation to nitric oxide neurotoxicity by PSD-95 protein. Science 284: 1845–8

Setou M, Nakagawa T, Seog DH, Hirokawa N (2000) Kinesin superfamily motor protein KIF17 and mLin-10 in NMDA receptor-containing vesicle transport. Science 288: 1796–802

Stenzel-Poore MP, Stevens SL, Xiong Z, Lessov NS, Harrington CA, Mori M, Meller R, Rosenzweig HL, Tobar E, Shaw TE, Chu X, Simon RP (2003) Effect of ischaemic preconditioning on genomic response to cerebral ischaemia: similarity to neuroprotective strategies in hibernation and hypoxia-tolerant states. Lancet 362: 1028–37

Stokin GB, Lillo C, Falzone TL, Brusch RG, Rockenstein E, Mount SL, Raman R, Davies P, Masliah E, Williams DS, Goldstein LSB (2005) Axonopathy and Transport Deficits Early in the Pathogenesis of Alzheimer’s Disease. Science 307: 1282–1288

Tanaka Y, Kanai Y, Okada Y, Nonaka S, Takeda S, Harada A, Hirokawa N (1998) Targeted Disruption of Mouse Conventional Kinesin Heavy Chain kif5B, Results in Abnormal Perinuclear Clustering of Mitochondria. Cell 93: 1147–1158

Terenzio M, Schiavo G, Fainzilber M (2017) Compartmentalized Signaling in Neurons: From Cell Biology to Neuroscience. Neuron 96: 667–679

Thomas GM, Huganir RL (2004) MAPK cascade signalling and synaptic plasticity. Nature Reviews Neuroscience 5: 173

Tjong YW, Jian K, Li M, Chen M, Gao TM, Fung ML (2007) Elevated endogenous nitric oxide increases Ca2+ flux via L-type Ca2+ channels by S-nitrosylation in rat hippocampal neurons during severe hypoxia and in vitro ischemia. Free Radic Biol Med 42: 52–63

Tovar KR, Westbrook GL (1999) The incorporation of NMDA receptors with a distinct subunit composition at nascent hippocampal synapses in vitro. J Neurosci 19: 4180–8

Tsien JZ, Chen DF, Gerber D, Tom C, Mercer EH, Anderson DJ, Mayford M, Kandel ER, Tonegawa S (1996) Subregion-and cell type-restricted gene knockout in mouse brain. Cell 87: 1317–1326

Tu W, Xu X, Peng L, Zhong X, Zhang W, Soundarapandian MM, Balel C, Wang M, Jia N, Zhang W, Lew F, Chan SL, Chen Y, Lu Y (2010) DAPK1 interaction with NMDA receptor NR2B subunits mediates brain damage in stroke. Cell 140: 222–34

Venere M, Horbinski C, Crish JF, Jin X, Vasanji A, Major J, Burrows AC, Chang C, Prokop J, Wu Q, Sims PA, Canoll P, Summers MK, Rosenfeld SS, Rich JN (2015) The mitotic kinesin KIF11 is a driver of invasion, proliferation, and self-renewal in glioblastoma. Science Translational Medicine 7: 304ra143–304ra143

Verhey KJ, Kaul N, Soppina V (2011) Kinesin Assembly and Movement in Cells. Annual Review of Biophysics 40: 267–288

Volk L, Chiu SL, Sharma K, Huganir RL (2015) Glutamate synapses in human cognitive disorders. Annu Rev Neurosci 38: 127–49

Wang X, Schwarz TL (2009) The Mechanism of Ca2+-Dependent Regulation of Kinesin-Mediated Mitochondrial Motility. Cell 136: 163–174

Wang Z, Cui J, Wong WM, Li X, Xue W, Lin R, Wang J, Wang P, Tanner JA, Cheah KSE, Wu W, Huang J-D (2013) Kif5b controls the localization of myofibril components for their assembly and linkage to the myotendinous junctions. Development 140: 617–626

Wild AR, Jones S, Gibb AJ (2014a) Activity-dependent regulation of NMDA receptors in substantia nigra dopaminergic neurones. The Journal of Physiology 592: 653–668

Wild AR, Jones S, Gibb AJ (2014b) Activity-dependent regulation of NMDA receptors in substantia nigra dopaminergic neurones. J Physiol 592: 653–68

Wong YL, Rice SE (2010) Kinesin’s light chains inhibit the head- and microtubule-binding activity of its tail. Proc Natl Acad Sci U S A 107: 11781–6

Wozniak MJ, Melzer M, Dorner C, Haring H-U, Lammers R (2005) The novel protein KBP regulates mitochondria localization by interaction with a kinesin-like protein. BMC Cell Biology 6: 35

Yin X, Takei Y, Kido Mizuho A, Hirokawa N (2011) Molecular Motor KIF17 Is Fundamental for Memory and Learning via Differential Support of Synaptic NR2A/2B Levels. Neuron 70: 310–325

Yip YY, Pernigo S, Sanger A, Xu M, Parsons M, Steiner RA, Dodding MP (2016) The light chains of kinesin-1 are autoinhibited. Proceedings of the National Academy of Sciences 113: 2418–2423

Zhou Q, Sheng M (2013) NMDA receptors in nervous system diseases. Neuropharmacology 74: 69–75

